# A complex evolutionary history of genetic barriers to gene flow in hybridizing warblers

**DOI:** 10.1101/2022.11.14.516535

**Authors:** Silu Wang, Graham Coop

## Abstract

The *Setophaga* warbler sister species in western North America, *Setophaga occidentalis* and *S. townsendi* provide a rare opportunity to explore how the accumulation of genetic barriers to gene flow contributes to speciation. These sister species are at the very early stages of speciation, as their genome-wide divergence is limited, and they hybridize extensively upon secondary contact. However, there are several well-characterized loci involved in speciation. There is a plumage difference between the species controlled by mutations close to the agouti signaling protein (ASIP), which generates underdominance due to compromised hybrid territorial signaling. A second large region (~2Mb) on chromosome 5 encodes many mitochondrial-interacting genes and coevolves with the mitochondrial genome. Here we reconstructed the ancestral recombination graphs of these genomic regions relative to the genome-wide background to learn about the evolutionary history of these barriers. We find signatures of long-lasting barrier effects that lasted for ~190 K generations, and recent selective sweeps in chr5 mitonuclear genetic block in both species within the past 60 K generations ago. We further observed heterogeneity within the long-lasting mitonuclear barrier with localized signals of much more recent selective sweeps unique one of the species. The divergence of ASIP is elevated by a species-specific selective sweep within *S. occidentalis* that occurred 21 K generations ago. However, the signature of long-lasting barrier effect is much weaker at ASIP suggesting that it may have arisen later than the mitonuclear system. Associating the evolutionary history of these genetic barriers with Pleistocene climate change history sheds light on the intrinsic and extrinsic reciprocity underlying the origin of species.

## Introduction

A major focus of evolutionary biology is on understanding how the accumulation of genetic barriers to gene flow contributes to reproductive isolation underlying the process of speciation. Key to this goal is understanding the relative timing that various types of barriers arise during speciation. For example, to what extent do differences in ecology, environments, or sexual selection pressures drive initial population divergence. There can also be genetic constraints on speciation that are expected to affect the timing of various types of barriers. Dobzhansky Muller Incompatibilities (DMIs, Dobzhansky 1937; Muller 1942) require changes at multiple loci and so can take time to accumulate (Orr and Turelli 2001). Furthermore, if hybridization occurs during the early stages of speciation, the resulting mendelian segregation and recombination can break down co-adaptive genetic combinations, such as DMIs (Slatkin 1975; Felsenstein 1981; Barton 1983; Butlin et al. 2021; MacPherson et al. 2022). Therefore, the accumulation of single-locus genetic barriers due to local adaptation or other divergent selection, could seed genomic differentiation with pairwise and higher order genetic barriers that only become prevalent later in speciation after the barriers to gene flow begin to congeal together (Franklin & Lewontin, 1970; Hartl 1977; Barton and Bengtsson 1986; Yeaman and Whitlock 2011; Feder et al. 2014). While there is a rich body of theory in this area, ultimately this is an empirical question, which recent work on the types of loci underlying reproductive isolation is starting to be addressed. This effort can be aided by reconstructing the divergence history of genetic barrier loci to estimate the order in which they spread.

The New World Wood Warbler sister species, *Setophaga townsendi* (STOW) and *S. occidentalis* (SOCC), are great systems for exploring the accumulation of barriers to gene flow (Fig. 1 A). This pair of sister species are at an early stage of speciation with a low level of genome-wide differentiation (FST ~ 0.03) (Wang et al. 2021). The species hybridize via stable hybrid zones (Fig. 1 A) as well as in ancient hybrid populations formed around 5000 years ago due to hybridization after the last glacial maxima (Fig. 1 A).

**Fig. 1.**
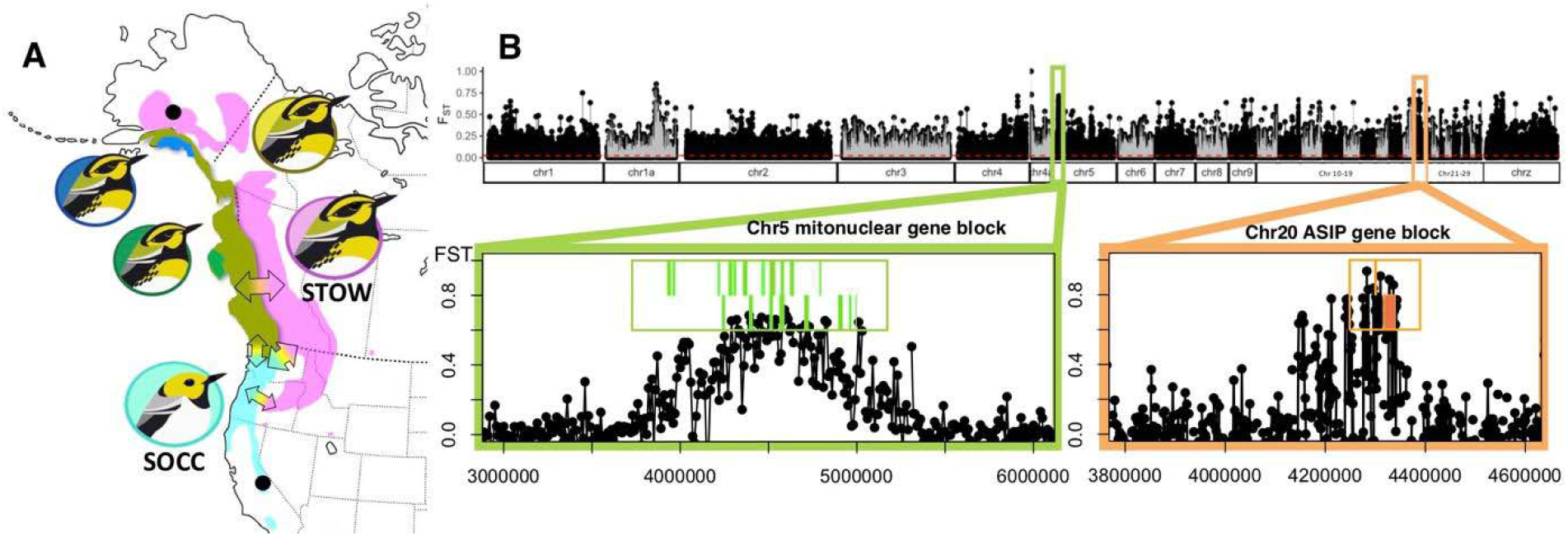
Distribution of ancestry of the SOCC-STOW species complex. **A**, the range of parental populations, SOCC and inland STOW, are respectively colored in turquoise and magenta. The ranges of ancient hybrid populations are colored in royal blue, ochre, and forest green. Arrows show regions of hybridization among the populations. Allopatric parental species sampling locations shown as black dots. **B**, F_ST_ scan along the genome with a zoom-in on the chr5 mitonuclear gene block and chr20 color gene blocks. Positions of genes on the forward strand (top) and reverse strand (bottom) are annotated within the boxed focal gene regions.

Several genomic targets of divergent selection have been recently characterized between these species using population genomic analyses of differentiation and hybrid zones clines (Fig. 1B). The first is a color polymorphism in the ASIP gene block that determines the differences of carotenoid and melanic plumage traits between species (Wang et al. 2020; de Zwaan et al. 2022). This appears to be involved in social signaling within species and may be maintained via underdominance due to conflicting signals in the heterozygote hybrid (Wang et al. 2020; de Zwaan et al. 2022).

The second set of loci are involved in an apparent mitonuclear incompatibility. The autosomal alleles in this system occur in a highly differentiated region on chromosome 5 that contains a number of genes involved in mitochondrial fatty acid metabolism (Fig. 1B). The two haplotypes at this chromosome 5 region show a strong statistically-association with two divergent mtDNA alleles among hybrids consistent with epistatic selection maintaining species-specific mitonuclear combinations, with signatures of climatic adaptation (Wang et al. 2021).

This warbler system with layers of hybridization and genomic barriers to gene flow identified (Fig. 1) provides a rare opportunity to unravel the accumulation of genomic barriers underlying speciation. To study the history of these loci we reconstruct coalescent genealogies along the genome using genome-wide sequence data from allopatric populations. Thanks to technical advancement in the reconstruction of ancestral recombination graphs we can now construct these genealogies for recombining sequences to more accurately learn about the history of speciation loci (Hejase et al. 2020; Campagna et al. 2022). This study elucidates the tension of selection and recombination in the early stage of speciation.

To understand the tension of selection and recombination in the early stage of speciation, here we reconstructed ancestral recombination graph of the genetic barriers and the randomly sampled genomic regions, and ask whether the genealogies at the color and mitonuclear loci (1) display signatures consistent with them being long-term ‘barriers to gene flow’; (2) show signals of species-specific selective sweeps (Fig. 2); (3) allow us to reconstruct their relative timing of divergence in the course of speciation.

**Fig. 2.**
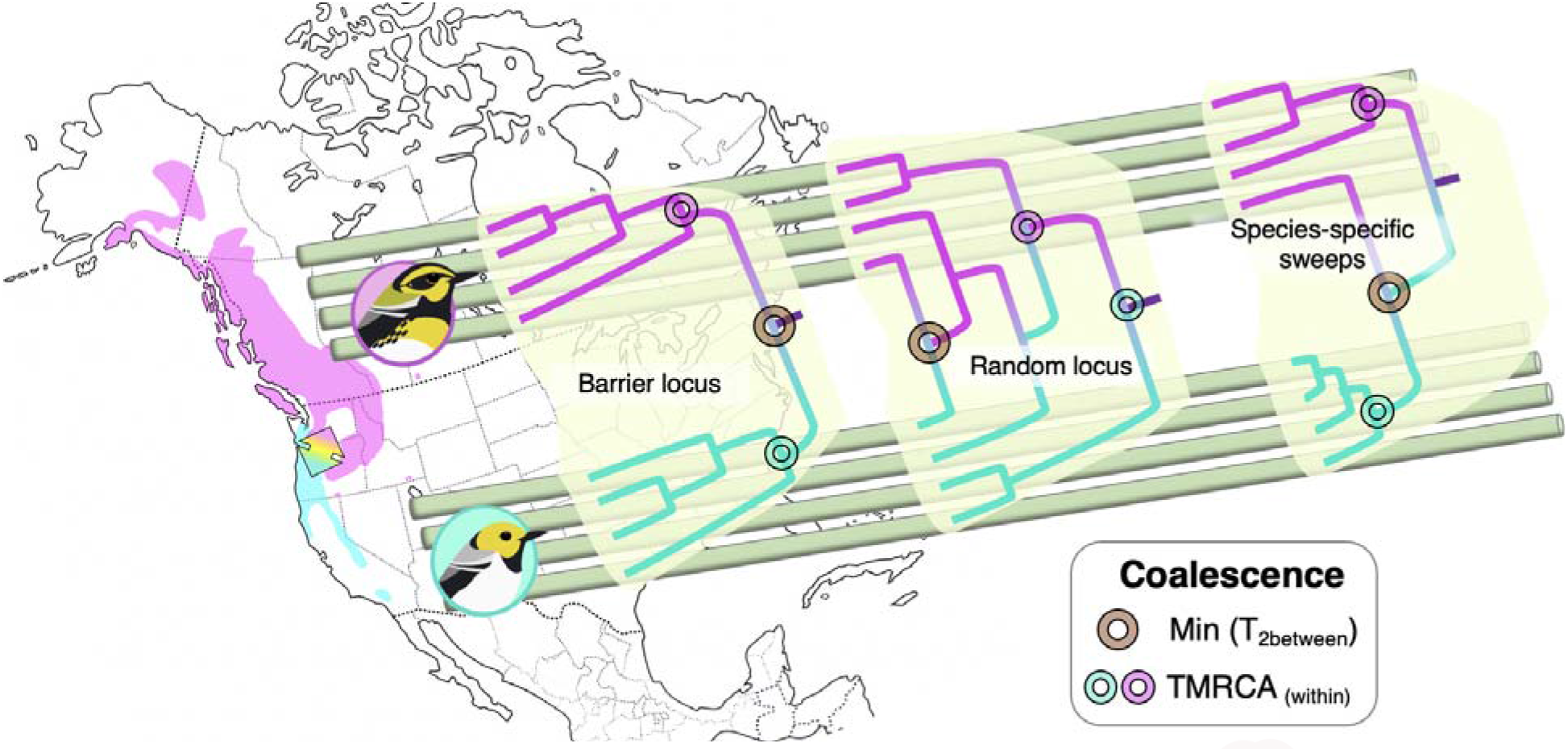
Illustration showing expected topology of coalescence trees of **(1)** a barrier locus, **(2)** a random locus, and **(3)** a locus under species-specific selection. **(1)** and **(3)** are not necessarily mutually exclusive at a genetic locus. Three different coalescent times are indicated on the trees: the minimum coalescent time between species (brown circle) and the TMRCA within SOCC and STOW (turquoise and magenta circles). See also Hejase et al. 2020.

### Expectations

Given the low levels of genome-wide differentiation between species, we expect genealogies in randomly chosen regions to show deep coalescent times within populations and extensive paraphyly (Fig. 2, random locus) due to gene flow and little time for lineage-sorting since the onset of speciation. Regions of high allele frequency differentiation, identified by measures of relative differentiation (F_ST_), can be due to reduced genetic diversity within populations and/or higher absolute divergence between populations (Charlesworth 1998; Cruickshank and Hahn 2014). Thus, loci associated with species differences can have either of these patterns and so there are two broad predictions for their genealogies.

First, these loci can be associated with reduced diversity within populations because the alleles underlying the differences can have rapidly swept up in frequency resulting in a short star-like phylogeny within the populations (Maynard Smith and Haigh 1974). These sweep genealogies will tend to be monophyletic as there has been little time for migration between populations (Fig. 2, species-specific sweeps).

Second, if the alleles underlying species differences are selected against in the opposing environment or genetic background, this results in a lower effective migration rate at these loci, i.e., they are barriers to gene flow, which can result in fewer coalescent events between populations increasing the chance of monophyly (Fig. 2, barrier loci). The time over which these barrier alleles have persisted will result in increased pairwise divergence between populations, due to deeper coalescent times between populations (although this in practice may be difficult to identify, unless barriers have prevented gene flow for a long time) (Charlesworth and Charlesworth 1997).

In practice, an adaptive substitution in one lineage may result in both a species-specific sweep and a barrier to migration, for example, in the case of a locally adapted allele that rapidly sweeps up in frequency to become a spatially balanced polymorphism. However, the signals can also occur in isolation. For example, a species-specific sweep due to a beneficial allele that is conditionally neutral in the second population and simply not spread yet because of insufficient time for neutral migration. Conversely an allele that serves as a barrier to migration could have hypothetically drifted up in frequency, or simply be old enough that any trace of a sweep has eroded away.

## Results

We reconstructed ancestral recombination graphs from sequence data from 5 birds from each species sampled from allopatric parts of the range (Fig. 1 A). We focused our analysis on the broad regions surrounding the ASIP and mitonuclear speciation genes and random regions in the genome (Fig. 3–5). At each SNP, we extract the marginal tree for each of the 1200 MCMC iterations, discarding a burnin period of 600 iterations. We calculate various summary statistics of the coalescent times from these trees for each iteration. We take the average of a given statistic over the iterations at a SNP (the posterior mean) and average these posterior means in 10kb non-overlapping windows for outlier tests. To find the marginal tree corresponding to the peak of coalescent time statistics, we conducted a finer scan with 5kb non-overlapping windows across candidate regions on chr5 and chr20. We then identified the marginal tree at the locus that peaks within the 5kb peak window.

**Fig. 3.**
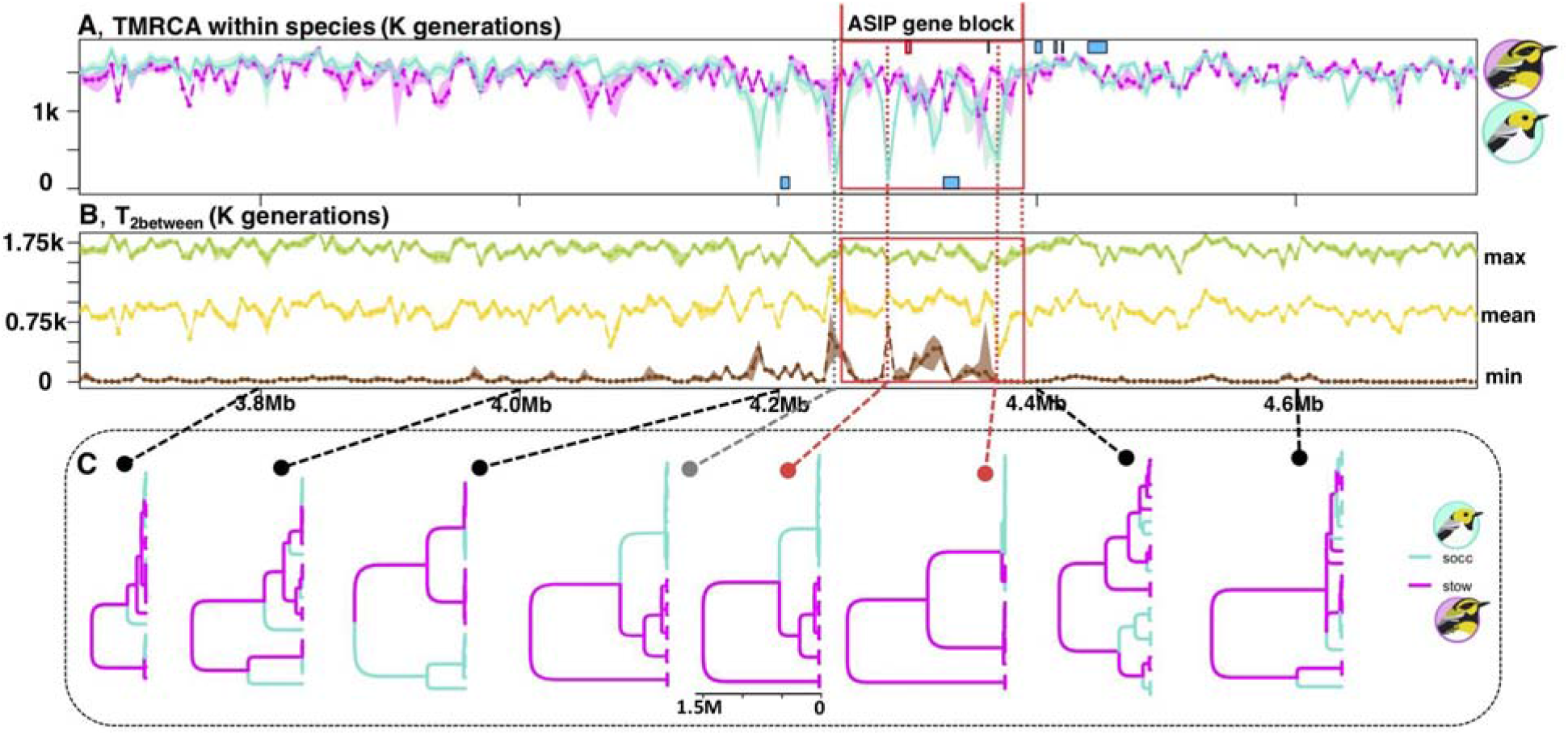
Coalescence history of ASIP and its flanking regions in chr20. **A**. The within-species TMRCA in SOCC (turquoise) and STOW (magenta). **B**, the minimum (brown), mean (gold), and maximum (green) pairwise coalescent times between species. All lines represent posterior means for 5kb windows, with the envelopes around the lines show the 95% credible intervals. **C**, examples of coalescent trees along genetic positions with branch lengths indicated by the generation scale bar. For better resolution of the coalescent tree topology, we find the 1kb window that peaks within each 5kb peak. There was a SOCC-specific selective sweep with TMRCA_socc_ being around 21 K generations ago. In contrast, TMRCA_stow_ was 1.48 M generations ago. All the genes in this region are indicated by the blue boxes with ASIP gene bordered by red and forward strand genes placed on the top and reverse strand genes on the bottom. There is an absence of genes in the further flanking regions of the ASIP gene block.

**Fig. 4.**
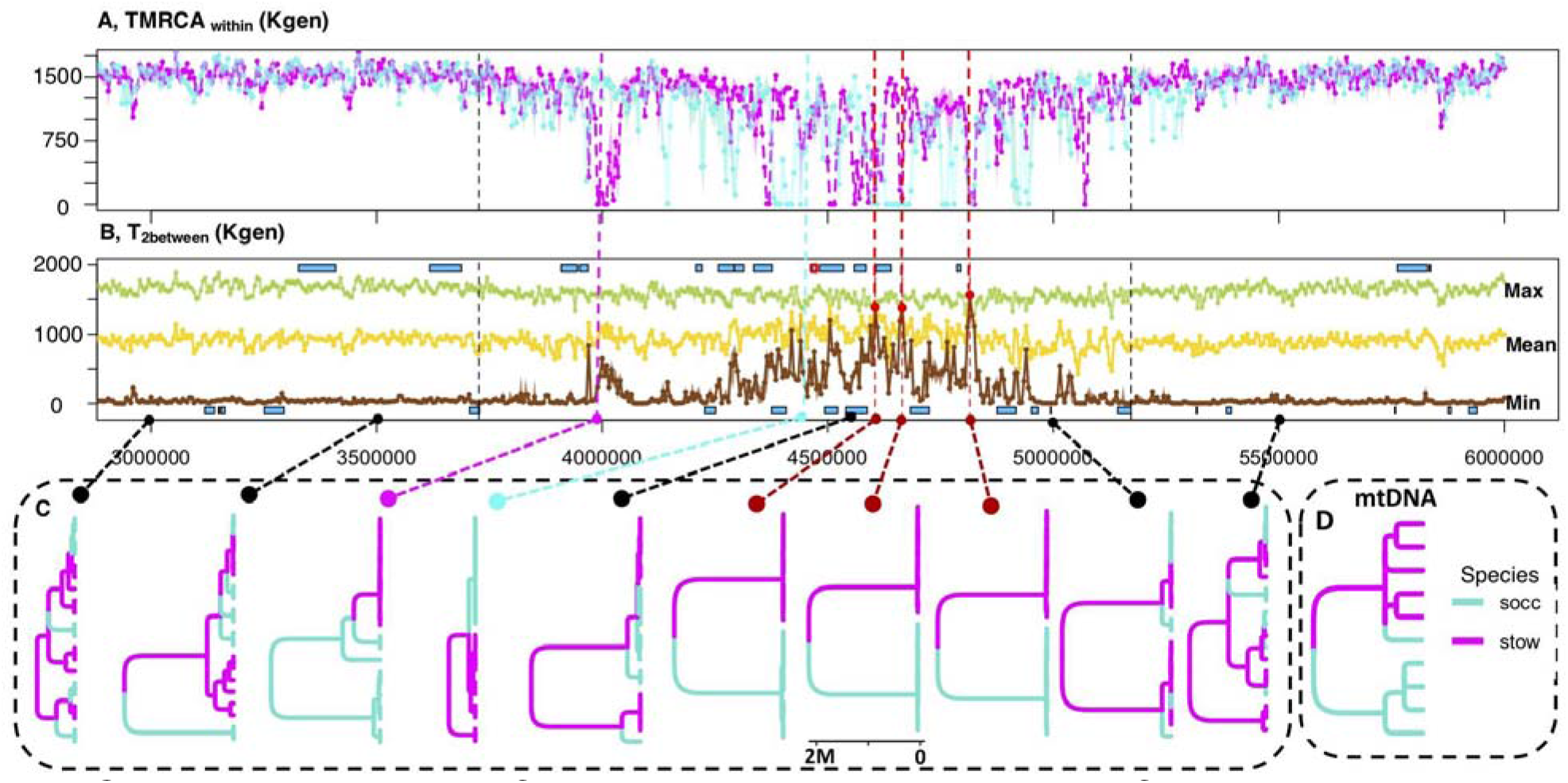
Coalescence history of chr5 mitonuclear genes and its flanking regions in chr5. **A**, The TMRCA within SOCC (turquoise) and STOW (magenta). Black dotted lines border the mitonuclear gene block. **B**, the maximum (green), mean (gold), and minimum (brown) pairwise between-species TMRCA. The blue rectangles indicate locations of protein coding genes in the forward (top) and reverse (bottom) strands. The gene with red border harbors the SNP that is correlated with mitochondrial haplotype in ancient hybrids (Wang et al. 2021). **C**, examples of coalescent trees of various genetic regions along genomic positions within and outside the mitonuclear gene block. Branch lengths are proportional to TMRCA as indicated by the scale bar. There are recent selective sweeps (red dots within the gene block) near *prostaglandin reductase 2* within both SOCC and STOW with TMRCA respectively 14 K (95%CI: 2.5-60 K) and 9.5 K (95%CI: 7-14.5 K) generations ago. There was also a STOW-specific sweep (TMRCA_STOW_ = 5 K generations, time discretization bounds: 2.5-7 K) within protein-coding gene *regulator G protein signaling 6* around 5 K generations. **D**. The coalescent tree of the mitochondrial genome with the same scale.

**Fig. 5.**
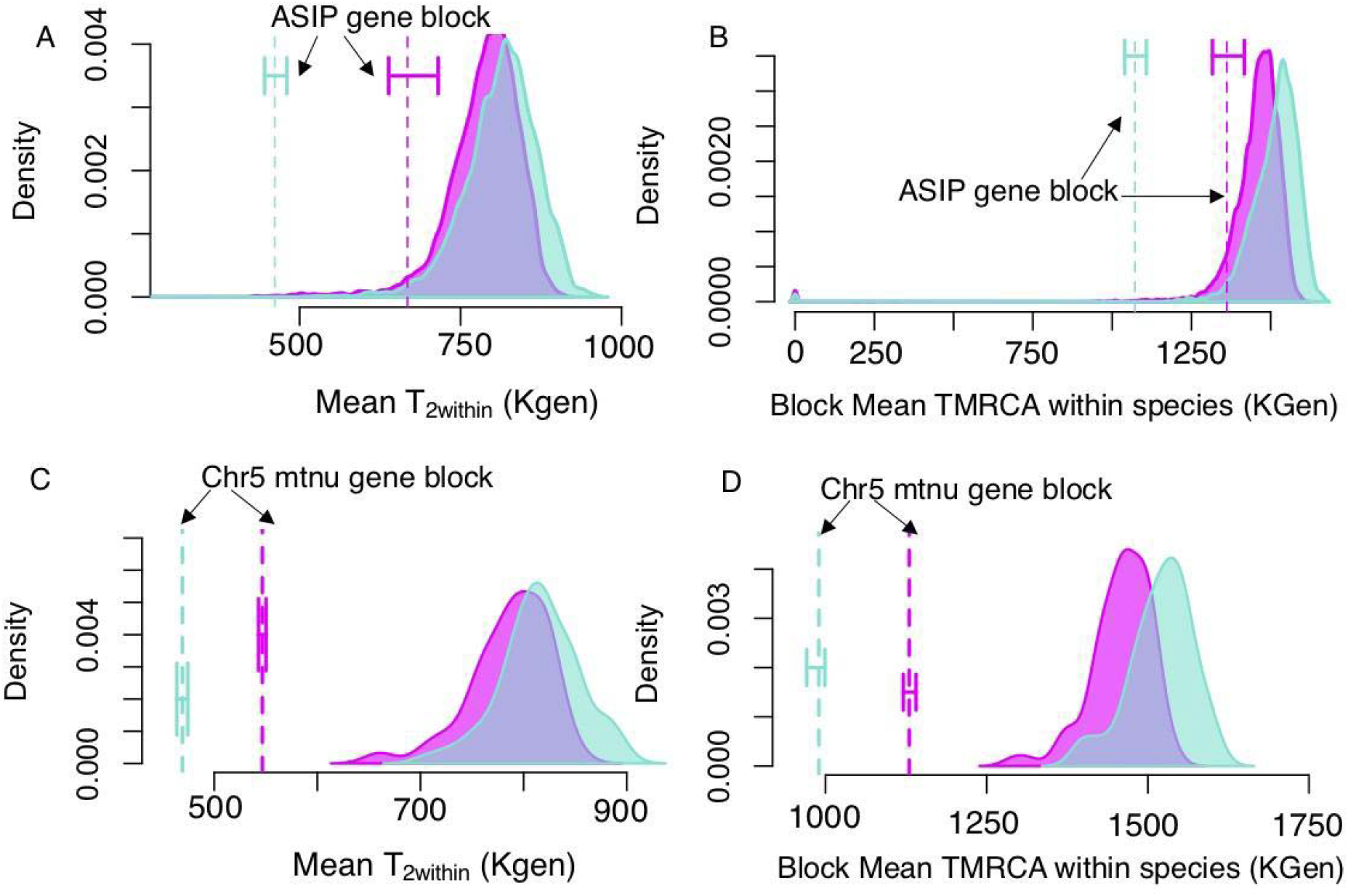
Genome-wide histograms of within-species (SOCC in turquoise and STOW in magenta) mean pairwise coalescent times (T_2within_) and TMRCA. **A** & **C**, Density plot of within-species mean **T_2within_** in randomly sampled gene blocks of the same size as the ASIP gene block (**A**) or the mitonuclear gene block (**C**) respectively. The vertical dotted lines are the mean T_2within_ of the ASIP gene block (**A**) or the mitonuclear gene block (**C**) with horizontal error bars representing the 95% posterior credibility interval. **B** & **D**, Density plot of the block mean within-species TMRCA in randomly sampled genetic blocks of the same size as the ASIP gene block (**B**) or the mitonuclear gene block (**D**) respectively. The vertical dotted lines indicate the mean of the candidate gene block, and the error bars are 95% credibility intervals.

In random regions of the genome the SOCC & STOW lineages are usually well mixed, as can be seen by the very recent minimum coalescent times between the species (e.g., in the flanking regions, brown lines Fig. 3 B and 4 B), indicating a high rate of gene flow between species over time. The deep genealogies in much of the genome stretching back to most common ancestor 1.5 M generations in the past on average, with similarly times to the most recent common ancestor within and between species (as in the flanking regions of Fig 3 & 4)

In contrast to these patterns, we found signatures of species-specific selective sweeps in both the ASIP color gene block (Fig. 3, 5) and mitonuclear gene blocks (Fig. 4–5). We further found the signature of long-standing barriers within the mitonuclear gene block, with a weaker signal in the ASIP gene block (Fig. 6).

**Fig. 6.**
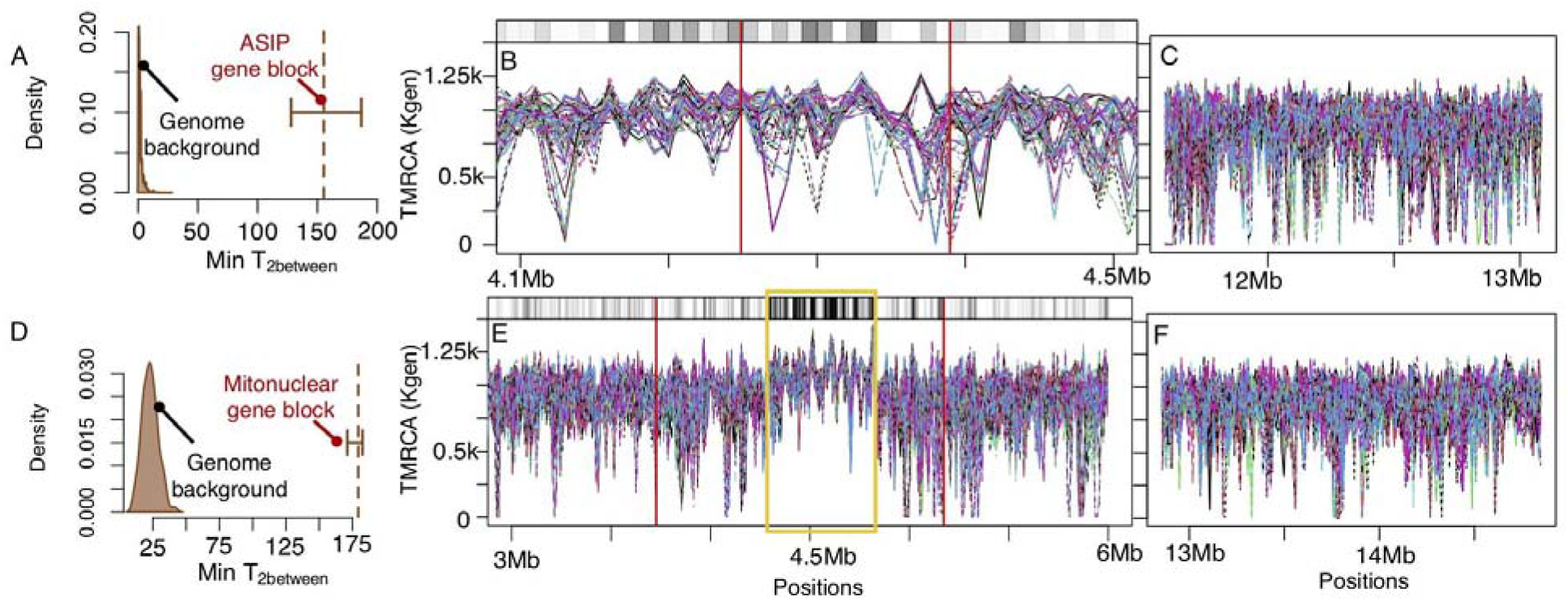
Signatures of long-standing barriers. **A & D**, the minimum T_2between_ is significantly greater in ASIP gene block (**A**) or the mitonuclear gene block (**D**) than the randomly sampled genomic blocks of the same size respectively. The dotted line represents the mean and error bars representing 95% CI among MCMC samples of the ARG estimates within the gene blocks. **B** & **E**, pairwise coalescent times of each between-species pair along ASIP gene block (**B**, bounded by vertical red lines) and mitonuclear gene block (**E**, bounded by vertical red lines) versus randomly sampled gene blocks of similar size (**C**, **F**). There is a long-standing barrier block (yellow box) within the chr5 gene block that is significantly diverged between species. The shaded bar of the top bar of **B** & **E** uses darker shade in 10kb bins to show the fraction of between-species pairwise coalescent times greater than 95% percentile of the genome.

### Species-specific sweeps: ASIP gene block

Looking across the broad ASIP region, the pairwise coalescent times within both species are comparable but the coalescent time times dip dramatically within SOCC in the ASIP gene block but not for STOW (Fig. 3 A). The marginal coalescent trees across this region (Fig. 3 C) show the trees adjacent to ASIP show a shallow monophyletic tree of the SOCC lineages and a much deeper paraphyletic tree in STOW. This is consistent with a recent species-specific sweep in SOCC (Fig. 3 A, C; Fig. 5 A-B).

Both the mean pairwise coalescent time (T_2socc_, Fig. 5 A) and time to the most recent common ancestor (TMRCA_socc_, Fig. 5 B) in SOCC are much lower within the ASIP gene block than randomly sampled equal-sized gene blocks from the rest of the genome. In contrast, the mean pairwise coalescent time is STOW (T_2stow_) and TMRCA_stow_ of the ASIP gene block did not significantly deviate from the rest of the genome (Fig. 5 A-B).

To estimate the approximate date of the SOCC-specific sweep within the ASIP gene block, we took the marginal tree from a position that is the closest to the TMRCA_socc_ trough among the 1kb windows within the 10kb trough of TMRCA_socc_. This yields an estimated sweep time of ~21 K (bounds: 15-21 K) generations in SOCC, while TMRCA_stow_ was much deeper (~1475 K, 95% CI = 1474 to 2103 K) (Fig. 3 C).

The minimum pairwise coalescent time between species, indicative of the time when most recent between-species migration occurred in the history of the sample, is elevated across the ASIP region (brown line Fig. 3 B; Fig. 6 A). This is consistent with alleles in the ASIP region forming a barrier to gene flow, but this interpretation is complicated by the signal of species-specific sweeps, and we return to this point in detail below.

### Species-specific sweeps: mitonuclear gene block

We found signals of recent species-specific sweeps within the broad 1.4Mb chr5 mitonuclear gene block in both SOCC and STOW (Fig. 4, 5 C-D). In this region, lineages of both species are often reciprocally monophyletic, coalescing rapidly to form a shallow genealogy within both species, with reasonable average deeper coalescent times between species (Fig. 4 C). The within-species entire block mean TMRCA were significantly more recent within the chr5 mitonuclear gene block than randomly sampled regions of similar size (Fig. 5 C-D). Within the mitonuclear gene block, the mean TMRCA_STOW_ is 1130 K (95% CI: 1120 to 1140 K) (Fig. 4 A, C) and TMRCA_socc_ is 990 K (95% CI: 970 to 1000 K). None of our randomly sampled regions had a TMRCA_stow_ or TMRCA_socc_ more recent than the mitonuclear gene block (Fig. 5 C-D).

We observe a heterogeneity of patterns within the broad chr5 gene block (Fig. 4 A). There were recent species-specific sweeps within this region with divergent haplotypes driven to high frequencies in the two species. The strongest divergently selected species-specific sweeps (see coalescence trees with red pointers) are recent (Fig. 4 C), with the TMRCA_socc_ and TMRCA_stow_ being respectively 14 K (95% CI: 2.5-60K) and 9.5 K (95% CI: 7-14.5 K). However, there are a number of other strong localized dips in between-species gene flow (Fig. 4 B), with reciprocal monophyletic trees (Fig. 4 C), which is possibly consistent with a number of sweeps occurring in this broader region in both species (Fig. 4 A). Therefore, the core chr5 region may have a complicated history of multiple relatively recent sweeps in both species. In addition, we also observed a somewhat separated STOW-specific selective sweep (2) near protein-coding gene regulator G protein signaling 6 (RGS6) (Fig. 4 C, magenta pointer) dated around 5 K generations (bounds: 3.5-7 K). Partitioning the chromosome 5 analysis by SOCC vs STOW associated mitochondrial haplotypes the within species minimum coalescent time between peaks within PAPLN (Fig. S1), which is in agreement with the previous mitonuclear GWAS which identified this autosomal gene as adjacent to the strongest association with mitochondrial type in a set of ancient hybrid populations (Wang et al. 2021).

### Long-standing barriers to gene flow

Both the ASIP and mitonuclear regions potentially show signals of being barriers-to-gene-flow, with deeper minimum coalescent times between species than their flanking regions (min T_2between_, Fig. 3 B, 4 B) and the genome background (Fig. 6). However, the deeper between-species minimum coalescent times can be a byproduct of species-specific sweeps as opposed to long-standing barriers to gene flow. This is because the rapid coalescence during species-specific sweeps results in fewer ancestral lineages older than the sweep, and so there are fewer opportunities to detect migrant lineages between the species, resulting in a deeper minimum coalescent time between the species (an issue also discussed by Hejase 2020, see their supplement). To robustly parse out this effect, we focus on pairs of lineages between the species; reasoning that by looking at a single lineage from each species we avoid the effects of the reduced sample of ancestral lineages within a species.

We traced the coalescent times of all the between-species pairs of lineages within the focal gene blocks and the genome background (Fig. 6). The reduced number of ancestral lineages in the swept regions can be seen directly in the coalescent trees (Fig. 3 C & 4 C) and in the lower variance in the pairwise between-species coalescent times as these represent a smaller number of independent events. If the focal gene block has been a long-standing barrier between species, the mean between-species TMRCA of this gene block should be greater than that of the randomly sampled equal-sized gene blocks (Charlesworth et al. 1997). In 10kb windows across our regions we examine the proportion of pairwise between-species coalescent times that are greater than the 95% cutoff for pairs elsewhere in the genome (a statistic whose expectation is not altered by the within species sweeps). Particularly in the chr5 mitonuclear region we see a broad region where the between species pair coalescent times are unusually deep. The average between-species coalescent time is 0.95 M generations in this chr5 region and 0.88 M in the rest of the genome, suggesting that a barrier to gene flow has persisted for ~70 K, generations on average in this region of the genome. Looking in the center in the region of significantly elevated pairwise differences (black bar bounded by yellow box Fig. 6 E), the between-species haplotype pair coalescence time was 1.07 M generations. The core long-standing barrier in this gene block lasted for 190 K generations.

## Discussion

We used ancestral recombination graphs to uncover a refined picture of the evolutionary processes underlying the early stages of speciation. We observed heterogeneity in the timing of selective sweeps and the signature of long-lasting barriers to gene flow within the divergent gene blocks. In particular, the mitonuclear gene block harbors long-lasting barriers to gene flow as well as recent species-specific sweeps at various time points in the recent past. In contrast, the color gene block exhibits weak signature of long-lasting barrier and recent SOCC-specific selective sweep.

The inferences of times come with the caveat that the scaling from genetic divergence to generations depends linearly on the mutation rate. Given that the mutation rate is extrapolated from other species, the absolute timing should be treated with caution. Therefore, we warn against drawing strong conclusions on the exact timing of the selective sweeps and the origin of barriers-to-gene-flow. However, the relative timings of events, e.g., the prolonged time gap between the formation of barriers and recent selective sweeps, should be much more robust to our uncertainty in the mutation rate.

### Pigmentation gene block

Changes in the ASIP gene block underpins species-specific melanic and carotenoid plumage pigmentation (Wang et al. 2020). Our analysis shows that a recent species-specific sweep occurred in SOCC but not in STOW, suggesting that SOCC has a derived, selected phenotype associated with this region (Fig. 3, 5 A-B). Consistent with this SOCC is the only species with clear-faced male breeding plumage among the closely related *Setophaga* species that diverged during the Pleistocene. A selective advantage of the loss of melanin on SOCC faces could be related to the thermoregulation role of plumage coloration, which influence the metabolic costs of homeostasis (Rogalla et al. 2022). SOCC breeds in southern Oregon and California, and so lighter plumage could be favored to reduce solar heating in its warmer breeding habitat compared to most *Setophaga* warblers. The SOCC-specific selective sweep within the ASIP gene block is dated to be around 37 Kya (at 1.8 years/generation, Milá et al. 2007), an interglacial period, suggesting that this difference has persisted for some time.

Although we only observed a weak signature of long-standing barrier effects of ASIP from coalescent genealogies, previous studies showed that it is currently a barrier to gene flow with steep, stable geographical clines in the hybrid zones (Wang et al. 2019, 2020). Thus, there can be a disconnect between the signal of long-term barriers to gene flow and their impact in the current day, perhaps because the ASIP barrier is relatively young. The selection against gene flow at the ASIP locus is at least partly due to the compromised territorial performance in hybrids with heterozygous ASIP genotypes due to divergence in territorial signaling between the species (de Zwaan et al. 2022). There could be additional selection against hybrids related to thermal-adaptation as indicated by the recent SOCC-specific selective sweep. Future studies could jointly investigate the social and thermal aspects selection at ASIP locus.

### Mitonuclear gene block

The divergence between the species in the chr5 mitonuclear gene block between the inland *S. townsendi* and *S. occidentalis* is old (with a divergence time of 1-2.1 M generations) as is the divergence of the species-associated mitochondrial haplotypes (with a divergence time of 1.1-1.4 M generations). Although the rest of the genome harbors extensive recent mixing between species, this region remains particularly divergently sorted and the core region potentially having resisted gene flow for 190 K generations between the allopatric populations.

Because of the relatively similar divergence periods of the chr5 mitonuclear gene block and mitochondrial genome, we cannot make a strong inference on the order of the events. There is no strong signal of a recent sweep on the mitochondrial of either species, with reasonable levels of mitochondrial diversity within both species (Wang et al. 2021, Fig. 4 D). However, there is good evidence of recent selective sweeps within the chr5 mitonuclear gene block in both species. This suggests that strong, ongoing divergent selection in both species on the autosomal side of mitonuclear coadaptation. This relatively gene-rich region, ~20 genes in around 2Mb, may have synergized multiple targets and forms of selection in the early stage of speciation in the face of gene flow. Several genes in this region are directly associated with mitochondrial function. Wang et al (2021) previously found that mtDNA clades and chr5 haplotypes strongly covary geographically in the old hybrid coastal STOW, consistent with epistatic selection. The autosomal gene, prostaglandin reductase 2 (PTGR2) is at the center of the long-standing barrier region (Fig. 4), thus it is a candidate gene of the initial divergence. PTGR2 is NADH-dependent and thus depend on mitochondrial gene product to function, and divergence at it could epistatically interact with genes such as the mitochondrial ND2, which has four non-synonymous substitutions between the divergent haplogroups (Wang et al. 2021).

The long-lasting barriers could shelter the spread of species-specific selective sweeps in this mitonuclear gene block in the face of ongoing gene flow. We have evidence of at least one sweep in each species and likely more. The recent STOW-specific selective sweep is dated around 9 Kya consistent with the timing of Douglas fir forests invasion along the Pacific Northwest after the last glacial maximum (LGM) (Pielou 1991). Climate change since the LGM could have driven strong selection on fat metabolism, dispersibility, and seasonal migration (Zink and Gardner 2017). The strongest signals of selective sweeps in both species and long-lasting barrier effect are again near PTGR2, which primarily regulates the lipid metabolism to maintain homeostasis. This gene and its interaction with the mtDNA in lipid metabolism could have been a target for divergent climate-related selection for homeostatic adaptation during prehistoric climatic oscillations, especially in the post-LGM warming around 20 thousand years ago. This could explain the covariation of local climatic variation and the ancestry of this mitonuclear gene block (Wang et al. 2021).

Somewhat further away in the same region there is a separate STOW-specific sweep near Regulator G protein Signaling (RGS6). RGS6 encodes for GTPase-accelerating protein, which regulates motor movement, cardiac functions, and night vision (Maity et al. 2012; Huang et al. 2018; Malerba et al. 2019). STOW has much greater migratory distance, and so variation at this gene could have contributed to migratory adaptation via one or more of these phenotypes. Future GWAS on migratory variation within STOW and in the hybrid zone would further validate this possibility.

### Speciation

This study only reveals part of the speciation process. There are other outlying regions of genomic divergence that many have contributed to the speciation process in this system. However, the color gene block and mitonuclear gene block represent the most outstanding genomic divergence genome-wide, thus our coalescent dissection of these genomic regions is a milestone for understanding speciation in this species complex. We observed quite variable timing to the evolutionary events underlying the speciation process; with the divergence of the ASIP gene block being much more recent than the long-standing barrier in the chr5 gene block and mitochondria. This suggests that the various steps of speciation have been quite protracted. This lag between different events could reflect the Pleistocene glacial cycles, which have driven dramatic environmental fluctuations and would have strongly influenced the range overlap of the two species.

The evolution of barriers-to-gene-flow may have begun from an initial seeding barrier in the chr5 gene block (Fig. 6 E, yellow box). The TMRCA in the chr5 mitonuclear gene block (Fig. 6 E) represents very old divergence between the species (~1-2.1 M generations) and the core of the region (Fig. 6 E) is estimated to have withstood gene flow for ~190 K generations. This may have sheltered the buildup of additional genomic divergence nearby, due to the reduced effective migration rate at linked loci supporting divergent selection or species-specific selection at nearby genes in the face of gene flow (Franklin & Lewontin, 1970; Hartl 1977; Barton and Bengtsson 1986; Feder et al. 2014). For example, the STOW-specific selective sweep close to RGS6 is ~0.6Mb from the core (Fig. 4) contributes to genetic divergence, but occurred much later than the initial divergence, potentially contributing to additional phenotypic divergence. Further work unpacking the phenotypes tied up in this region could reveal how ancient barriers facilitate the establishment of the more recent genetic divergence nearby. Altogether the barriers in the mitonuclear gene block on chr5, the mitochondrial genome, and the color gene block on chr20 illustrate the genetic progression of speciation over tens of thousands of years.

## Acknowledgements

This work was supported by funding provided by the National Institutes of Health (NIH R35 GM136290 awarded to GC).

---

**Fig. S1.**
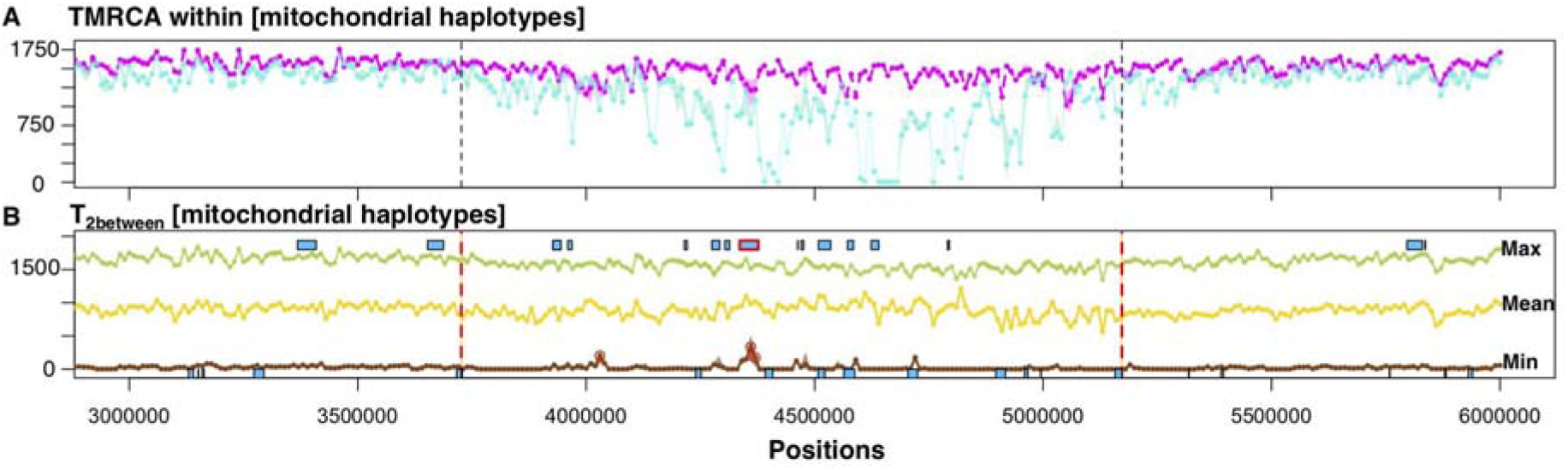
Coalescence time of SOCC (turquoise) versus inland STOW (magenta) mitochondrial haplotypes. **A**, TMRCA within haplotypes. **B**, Max (green), Mean (gold), and Minimum (brown) coalescent time between haplotypes. Blue boxes represent locations of genes with the red-bordered gene contains the SNP that covaries with mitochondrial haplotypes in ancient hybrids (Wang et al. 2021).

## Methods

### Whole genomic sequence data

We analyzed the published whole genome sequencing data from 5 individuals sampled from the allopatric range of *Setophaga townsendi* and *Setophaga occidentalis* (previously reported in Wang et al. 2020; Baiz et al. 2021). We trimmed and aligned the reads to the *Setophaga coronata* reference (version 2.1) (Baiz et al. 2021), identified variable sites following the pipeline given in Wang et al. 2021.

### Ancestral recombination graph

To reconstruct the evolutionary history of genetic barriers and the rest of the genome. We reconstructed the full history of recombination and coalescence with ARGweaver (Rasmussen et al. 2014). We executed 1200 iterations of MCMC runs with arg-sample command with the following priors: mutation rate = 4.42×10^−9^ (Nam et al. 2010), recombination rate = 1.5×10^−8^, effective population size = 452897, as π/4μ = 0.0040/(4×4.42×10^−9^). We specified unphased genomic data for ARGweaver to search and integrate over all the phasing options. The mitochondrial divergence was previously investigated by Wang et al (2021). To facilitate comparisons here we also ran ARGweaver for the mitochondrial genome specifying a recombination rate of zero and a mutation rate used in Wang et al (2021) of 2×10^−8^. We masked the missing data with the *maskmap* command. To generate the genomic map of missing sites, we used bedtools. To ensure haplotype-calling quality, focused on sites with a minimum vcf quality of 30 (different choices of quality cutoff did not substantially alter the findings). To allow sufficient time resolution for coalescence events that happened in the distant past, we included more time points with ntimes = 30, as opposed to the default time points of 20. We set the maximum coalescence time to be 3 million (M) generations as slightly more than twice the estimated value of 4Ne. We kept the default setting for the rest of the commands.

To track the local coalescence genealogies among individuals within and between species, we monitored coalescence tree statistics along genomic positions. For each coalescence tree calculated from each genomic region in each iteration, we calculated the minimum, mean, and maximum of pairwise between-species (T_2between_) and within-species (T_2within_) coalescence time (Fig. 1B). We take the mean of the variable within 10kb of non-overlapping windows for outlier tests. To visualize the coalescence timing within and around the candidate regions of interest, we plot the 5kb non-overlapping windows. To find the coalescent trees associated with minimum T2between peaks, we identified the coalescent tree that is closest to the 1kb window that peaks within the 10kb peak. We estimated 95% credible intervals of TMRCAs from the post-burn iterations of the MCMC. As ARGweaver works with discretized times, if the TMRCA of interest did not vary substantially across the MCMC we quote the discretized intervals that bracket the TMRCA as “bounds” on the time in the main text.

The sequence processing and analytical pipeline in this study is deposited here https://github.com/setophaga/Speciation.gene.ARG .

### Gene region boundaries and genomic background

The ASIP gene block is delineated as the ASIP and its neighboring genes within 50 Kb flanking regions on both sides. The mitonuclear gene block is delineated by FST peaks and LD elevation in ancient hybrids (Wang et al. 2021). To understand coalescent history in the rest of the genome, we sampled 1.5 Mb of gene block starting at 3.5 Mb from both chromosomal ends (like the putative speciation genes on chr5 and chr20) and a randomly selected gene block within each chromosome. We compared the mean test statistics within the ASIP and mitonuclear gene block to the respective gene blocks of similar size in the genomic sampling.

